# Allogeneic Tumor Cell-Derived Extracellular Vesicles Stimulate CD8 T Cell Response in Colorectal Cancer

**DOI:** 10.1101/2023.04.17.537250

**Authors:** Travis J Gates, Dechen Wangmo, Xianda Zhao, Subbaya Subramanian

## Abstract

Colorectal Cancer (CRC) is the second leading cause of cancer-related death in the United States. Most CRC patients present with a microsatellite stable (MSS) phenotype and are highly resistant to immunotherapies. Tumor extracellular vesicles (TEVs), secreted by tumor cells, can contribute to intrinsic resistance to immunotherapy in CRC. We previously showed that autologous TEVs without functional miR-424 induce anti-tumor immune responses. We hypothesized that allogeneic modified CRC-TEVs without miR-424 (mouse homolog miR-322) derived from an MC38 background would effectively stimulate CD8^+^ T cell response and limit CT26 tumor growth. Here we show that prophylactic administration of MC38 TEVs without functional miR-424 significantly increased CD8^+^ T cells in CT26 CRC tumors and limited tumor growth, not B16-F10 melanoma tumors. We further show that the depletion of CD4^+^ and CD8^+^ T cells abolished the protective effects of MC38 TEVs without functional miR-424. We further show that TEVs can be taken up by DCs *in vitro,* and subsequent prophylactic administration of autologous DCs exposed to MC38 TEVs without functional miR-424 suppressed tumor growth and increased CD8^+^ T cells compared to MC38 wild-type TEVs exposed to DCs, in Balb/c mice bearing CT26 tumors. Notably, the modified EVs were well tolerated and did not increase cytokine expression in peripheral blood. These findings suggest that allogeneic-modified CRC-EVs without immune suppressive miR-424 can induce antitumor CD8^+^ T cell responses and limit tumor growth in vivo.

## INTRODUCTION

Colorectal cancer (CRC) is the third most common cancer diagnosis in the US and the second leading cause of cancer-related deaths, and the incidence is increasing in patients <50.^1^ Immune checkpoint inhibitors (ICIs) nivolumab (anti-PD1) and ipilimumab (anti-CTLA4) have drastically expanded treatment options for multiple malignancies of various tissue types.^2–4^ However, only CRC patients with microsatellite instability-high (MSI-H) subtype receive any clinical benefit to ICIs accounting for only <15% of CRC patients.^1, 5–8^ Most patients (∼85%) present with microsatellite stable (MSS) disease and are resistant to ICIs.^6, 9^ MSI-MSS stratification is used to predict the efficacy of ICIs in CRC.^6, 7, 10^ It has been demonstrated that immune cell infiltration is only observed in a small proportion of MSS-CRC.^11, 12^ There is a critical clinical need to determine intrinsic resistance mechanisms and increase tumor T cell infiltration,^13, 14^ to synergize with current ICIs to treat CRCs.

Tumor-immune cell crosstalk within the CRC tumor environment can influence the antitumor immune response mounted by CD8^+^T cells and efficacy to ICIs. Mechanisms of tumor-immune cell crosstalk mediated by tumor-secreted extracellular vesicles (TEVs) have been shown to influence the phenotype and function of immune cells in the TME.^15, 16^ TEVs can be immunosuppressive or immunogenic in a context-specific manner, depending on the cargo contained within TEVs. For instance, PD-L1 abundance in TEVs has yielded local immunosuppression.^17^ Further, depletion of PD-L1 expression in TEVs can reverse immunosuppressive effects.^18^ Our previous investigation also showed that syngeneic TEVs harboring miR-424 could reduce CD28/CD80 costimulatory gene expression and contribute to immune suppression and ICI resistance in CRC.^19^ Conversely, tumor antigens and other immunomodulatory agents such as IFNγ and dsDNA within TEVs can stimulate immune responses.^20–22^ Early experiments of TEV-immune regulation were done in a syngeneic context.^23^ However, modifying TEVs in a syngeneic context presents a challenge in translating into clinical implementation due to insufficient tumor biomass.^23^

Allogeneic EVs from dendritic cells (DCs) have been shown to produce effective antitumor immune responses and improved T cell memory.^24^ Other efforts have attempted to expose DCs to TEVs and other tumor antigens, which showed success in preclinical models.^25^ However, the clinical benefit of DC vaccine approaches is limited to Sipluecel-T in metastatic castration resistant prostate cancer.^26^ Pre-clinical models of DC vaccine usage in CRC has shown inconsistent efficacy results.^27^ TEV cargo containing miR-424 could contribute to local immune suppression even in the presence of tumor antigens by suppressing T cell costimulation.^19^ Here we assessed the effects of allogeneic MC38 colon cancer TEVs that lack functional miR-424 (MC38-424i). We show that prophylactic administration of MC38-424i TEVs significantly limited tumor growth in mice bearing CT26 colon cancer but not B16-F10 melanoma cells. Additionally, we show that miR-424i TEVs can be pulsed to DCs and autologously transferred to Balb/c mice, leading to increased CD8^+^T cells in tumors. Future investigations should assess how allogeneic modified TEVs pulsed to DCs can synergize with ICIs in CRC for clinical translation.

## MATERIALS AND METHODS

### Mice and Animal Husbandry

All animal studies were approved by the Institutional Animal Care and Use Committee (IACUC) of the University of Minnesota. All mice were housed in pathogen-free conditions with fully autoclaved cages to minimize non-tumor-specific immune activation. Balb/cj and C57BL/6j mice were purchased from the Jackson Laboratory. Mice were bred in-house and were used for experiments between 6-8 weeks of age.

### Cell Lines and Cell Culture

Mouse CRC cell line CT26 (ATCC CRL-2638) was purchased from ATCC. Mouse CRC cell line MC38 was kindly provided by Dr. Nicholas Haining. MC38 cells stably expressing miR322 inhibitor mouse homolog to miR-424 (MC38-424i), and MC38-miR-control cells were used in this study as described in ^19^. CT26 cells were cultured in complete Roswell Park Memorial Institute Medium (RPMI) 1640 medium (Gibco), supplemented with 10% heat-inactivated fetal bovine serum (FBS) (Thermo Fisher Scientific), 100 IU/mL penicillin, and 100 μg/mL streptomycin (Invitrogen Life Technologies). MC38 WT, MC38-424i, and MC38-miRcontrol cells were cultured in the complete Dulbecco’s modified Eagle medium (DMEM) (Gibco), supplemented with 10% heat-inactivated fetal bovine serum (FBS) (Thermo Fisher Scientific), 100 IU/mL penicillin, and 100 μg/mL streptomycin (Invitrogen Life Technologies). B16-F10 melanoma cells (ATCC CRL-6475) were purchased from ATCC and were cultured in the complete Dulbecco’s modified Eagle medium (DMEM) (Gibco), supplemented with 10% heat-inactivated fetal bovine serum (FBS) (Thermo Fisher Scientific), 100 IU/mL penicillin, and 100 μg/mL streptomycin (Invitrogen Life Technologies). Cell lines were authenticated and routinely tested for mycoplasma.

### Isolation of Tumor Extracellular Vesicles

24 hours prior to tumor extracellular vesicle (TEV) isolation, cell media was changed to DMEM media supplemented with 10% exosome depleted FBS (Gibco), 100 IU/mL penicillin, and 100 μg/mL streptomycin (Invitrogen Life Technologies). A standardized differential centrifugation protocol was used to purify TEVs from cell culture supernatants from MC38 WT, MC38-424i, and MC38 miR-control cells. Cell culture supernatants were centrifuged at 300g for 10 minutes to remove cells. Supernatants were centrifuged for 3,000g for 10 minutes to remove dead cells. Supernatants were centrifuged at 10,000g for 30 minutes to remove cellular debris. Supernatants were centrifuged at 2000g for 30 minutes in Amicon Ultra-15 (Millipore) to concentrate supernatant. Supernatants were ultracentrifuged at 100,000g for 70 minutes at 4LC with a SW40Ti rotor (Beckman Coulter). Pelleted TEVs were suspended and washed in PBS and underwent ultracentrifugation at 100,000g for 70m minutes at 4LC. TEVs were collected in PBS for downstream analysis and experimentation.

### Characterization of Tumor Extracellular Vesicles

To characterize the purified TEVs, we first used electron microscopy. TEVs suspended in PBS were placed on formvar carbon-coated nickel grids. TEVs on grids were stained with 2% uranyl acetate and allowed to air dry. TEVs were visualized using an FEI Tecnai G2 F30 Field Emission Gun Transmission Electron Microscope with a 4k x 4k ultrascan charge-coupled device camera. The size distributions and concentration of TEVs isolated from cell culture supernatants were determined using the NanoSight LM-10 microscope (Malvern Instruments) equipped with particle tracking software. 10 independent microscopic fields were captured and analyzed per cell line sample. Data were merged and presented as a single histogram plot. Additionally, we tested TEV related protein markers from isolated TEVs using western blotting. TEV protein concentration was estimated by using the Pierce Micro BCA Protein Assay Kit. 10ug of TEV protein from MC38 and modified cell lines were loaded on SDS gels. Primary antibodies binding markers: CD81 (cat# 104902, Biolegend 1:1000), and ALIX (cat# 634502, Biolegend 1:2000) were used to validate TEV protein markers. B-actin (cat# 8H10D10, Cell Signaling, 1:1500) and B-tubulin (cat# MA5 16308, Invitrogen,1:2000) confirmed no cellular contaminates.

### Prophylactic modified MC38 TEVs administration in CT26 and B16-F10 Subcutaneous Tumor Models

10µg of MC38 WT, MC38-424i, and MC38 miR-control TEVs suspended in sterile saline control were prophylactically injected twice into the tail vein of Balb/c or C57BL/6 mice depending on the experimental set up. We allowed two weeks for an adaptive immune response prior to CT26 or B16-F10 tumor cell challenge. To establish subcutaneous tumors, we injected 2×10^5^ CT26 colon cancer cells suspended in 100µL of 50:50 RPMI:Matrigel (Corning) into the right flank of Balb/c mice. Further, 2X10^5^ B16-F10 melanoma cells were injected in the same preparation into the right flank of C57BL/6 mice. After tumor cell inoculation, tumors were measured 3 times per week using an electronic caliper. Tumor volumes were calculated using the formula [volume = (width^2^ x length)/2]. Mice were sacrificed at 21 days following tumor inoculation. Tumor tissues were excised, imaged, and fixed in 10% neutral buffered formalin overnight. Fixed tissues were paraffin-embedded and sectioned in the University of Minnesota Clinical and Translational Sciences Institute. Mouse peripheral cytokines were measured using the Mouse Proteome Profiler Kit (R&D Systems) following the manufacturer’s protocol on 100uL of mouse serum from whole blood.

### Immunofluorescence of Tumor Tissues

Formalin fixed paraffin embedded (FFPE) tissues were deparaffinized with 3 xylene washes, rehydrated with gradient ethanol, and underwent antigen retrieval in antigen retrieval buffer (AR9, PerkinElmer) in a 95LC water bath. Sections were blocked in 5% bovine serum albumin buffer for 30 minutes. Primary anti-mouse antibody CD8 (cat# ab217344, Abcam, 1:100) was added and incubated overnight at 4LC. The tissue sections were washed twice with PBS and incubated with secondary antibodies (goat-anti-rat-A568, cat# A11077, Invitrogen, 1:250) and (goat-anti-rabbit-A568, cat# A11011, Invitrogen, 1:250) for 1 hour. Tissues were washed twice with PBS, and slides were mounted with slide mounting media with DAPI (cat# ab104139, Abcam). Slides were imaged on the BZX810 fluorescence microscope (Keyence). Quantitative image analysis was performed by counting positive signal percentage and fluorescence intensity signal in at least 5 randomly selected fields of each tumor tissue core.

### CD4-CD8 T cell Depletion

T cell subsets were depleted by administering 400µg *i.p.* of depleting antibody twice prior to prophylactic administration of MC38 TEVs and CT26 tumor challenge. CD4 T cells were depleted with anti-CD4 mAb (Clone GK1.5, BioXcell) CD8 T cells were depleted with anti-CD8α (Clone 2.43, BioXCell). CD4 and CD8 T cell depletion were confirmed using flow cytometry on the BD FACS CantoII (BD Biosciences) from the mouse spleen, lymph node, and peripheral blood. Antibodies for flow cytometry were CD3-APC (cat# 100236, Biolegend, 1:200) CD8-APCCy7 (cat# 100714, Biolegend, 1:100) CD4-BV510 (cat# 100449, Biolegend, 1:100) CD11b-FITC (cat# 101206, Biolegend, 1:100) CD19-FITC (cat# 115506, Biolegend, 1:100) Nk1.1-FITC (cat# 108706, Biolegend, 1:100) CD45-PE (cat#103106, Biolegend, 1:200). um

### Monocyte Isolation and Dendritic Cell Differentiation

Monocytes were isolated using a negative selection from mouse spleen, inguinal, axillary, and brachial lymph nodes with the Dynabeads Mouse DC Enrichment Kit (cat# 11429D, Invitrogen) according to the manufacture’s protocol. Monocytes were plated at 10^7^ cells/well and differentiated with Dendritic Cell Differentiation Kit (cat# CDK004, R&D Systems) according to the manufactures instructions. DCs were differentiated for 6 days in the kits’ media supplemented with 250IU/mL IL4 and 800 IU/mL granulocyte macrophage colony stimulating factor for two days. Then the cells were centrifuged and cultured in fresh complete media supplemented with 2,000 IU/mL IL4 and 2,000 IU/mL granulocyte macrophage colony stimulating factor. On day 6, cells were centrifuged and resuspended in media supplemented with 2,000 IU/mL IL6, 400 IU/mL IL1β, 2,000 IU/mL tumor necrosis factor α, and 100ng/mL lipopolysaccharide and were cultured for 24 more hours. On day 7, we confirmed DC differentiation with flow cytometry to see the percentage of cells expressing a high level of MHCII. Antibodies in flow cytometry were CD3-FITC (cat#100204, Biolegend, 1:100), CD19FITC (cat#115506, Biolegend, 1:100), Nk1.1-FITC (cat# 108706, Biolegend, 1:100) CD45-PE (cat# 103106, Biolegend, 1:200), CD11b-APC (cat# 101212, Biolegend, 1:100), IA/12E-APC-Cy7 (cat# 107628, Biolegend, 1:100) CD11c-BV510 (cat# 117338, Biolegend, 1:100)

### Dendritic Cell TEV-Uptake Experiment

DCs and TEVs were isolated as previously described. DCs were plated on Labtek II chambers with chamber protectors coated with fibronectin (5ug/mL) and differentiated in the same manner as previously described. TEVs from MC38 cell lines were stained with DiO lipophilic dye (Invitrogen) for 30 minutes and washed three times with PBS and ultracentrifugation at 100,000g for 70 minutes at 4LC. TEV pellets labelled with DiO were suspended in DC media, and 1µg TEVs were added to DC cultures on Day 6 with 100ng/mL LPS. Following 24 hours to allow for TEV uptake DCs culture on glass slides was stained with Cytopainter Red (cat# ab219942, Abcam) for 30 minutes according to the manufactures instructions and fixed with 4% formaldehyde for 30 minutes and washed 5 times with PBS. Chamber protectors were removed, slides were mounted with mounting media with DAPI, and DCs exposed to MC38 TEVs were imaged on the BZX810 fluorescence microscope (Keyence). A series of photos were taken with a (4.4 µm) stepwise increase (0.4 µm) at each step on the zaxis to validate the intracellular uptake of TEVs prior to the *in vivo* experiment.

### Prophylactic Autologous TEV-Dendritic Cell Animal Model

DCs were isolated and differentiated as previously described from Balb/c mice. TEVs from MC38 WT, MC38 322i, MC38 miRi-control were isolated as previously described. On day 6 10ug of TEVs from the MC38 cell lines were administered to DCs in culture. On day 7 DCs were scraped from 10cm^3^ dishes and counted and dosed at 1×10^6^ cells/injection in 100µL of sterile PBS. 5 groups (n=5/group) of animals were injected prophylactically with *i.v.* tail vein injection of autologous DC cell suspensions of (MC38 WT-DCs, MC38 322i-DCs, MC38 miRi-control-DCs, No TEV control-DCs, and saline. Balb/c mice underwent 2×10^5^ CT26 subcutaneous tumor challenge 14 days following the administration of DCs. Tumors were measured 3 times a week for 21 days using an electronic caliper. Mice were sacrificed at 21 days following tumor inoculation. Tumor tissues were excised, imaged, and fixed in 10% neutral buffered formalin. Fixed tissues were paraffin embedded and sectioned in the University of Minnesota Clinical and Translational Sciences Institute.

### Statistics

We used GraphPad Prism, versions 6.0 and 8.0, to perform statistical analyses and visualize data. We used the Student t-test to compare the treatment and control arms. One-way ANOVA was used when comparing more than 2 groups. All data are plotted as the mean ± Standard Error of the Mean (SEM). All statistics were evaluated at two-tailed α=0.05 unless otherwise corrected for multiple comparisons.

## RESULTS

### Allogeneic modified CRC-TEVs increase T cell infiltrates and decrease tumor growth

Previously we showed that colorectal cancer tumor cell-derived extracellular vesicles (CRC-TEVs) contained immunosuppressive miR-424 in a syngeneic context^19^; we sought to test if allogeneic modified TEVs (EVs lacking functional miR-424) derived from MC38 cell lines of a C57BL/6 background affected Balb/c mice bearing CT26 tumors. To test this, we used MC38 wild type (MC38 WT) colon cancer cell lines and modified cell lines that stably express miR-424i inhibitor (MC38-424i) and MC38-424i scramble control (MC38 miR-control). We first confirmed the quality of TEVs isolated from each cell line by western blotting for known EV markers CD81 (∼25kDa), and ALIX (∼100 kDa). We also included β-actin (∼40 kDa) and β-tubulin (∼60 kDa) to ensure no known contaminates in TEVs **Figure 1A**. We observed the expression of ALIX and CD81 in MC38 WT, MC38-424i, and MC38 miR-control TEVs, but did not observe the expression of cellular markers β-actin and β-tubulin. We then performed a Nanotracker analysis on MC38 TEVs to determine the size distribution of MC38 WT, MC38-424i, and MC38-miRcontrol TEVs **Figure 1B**. Average size distributions for TEVs were measured to have an x50 diameter of 136.9, 144.8, and 97.1nm for MC38 WT, MC38-424i, and MC38-miR-control TEV groups, respectively. We then performed Transmission electron microscope (TEM) imaging on the TEV and confirmed the morphology of TEVs **Figure 1C**.

**Figure 1).**
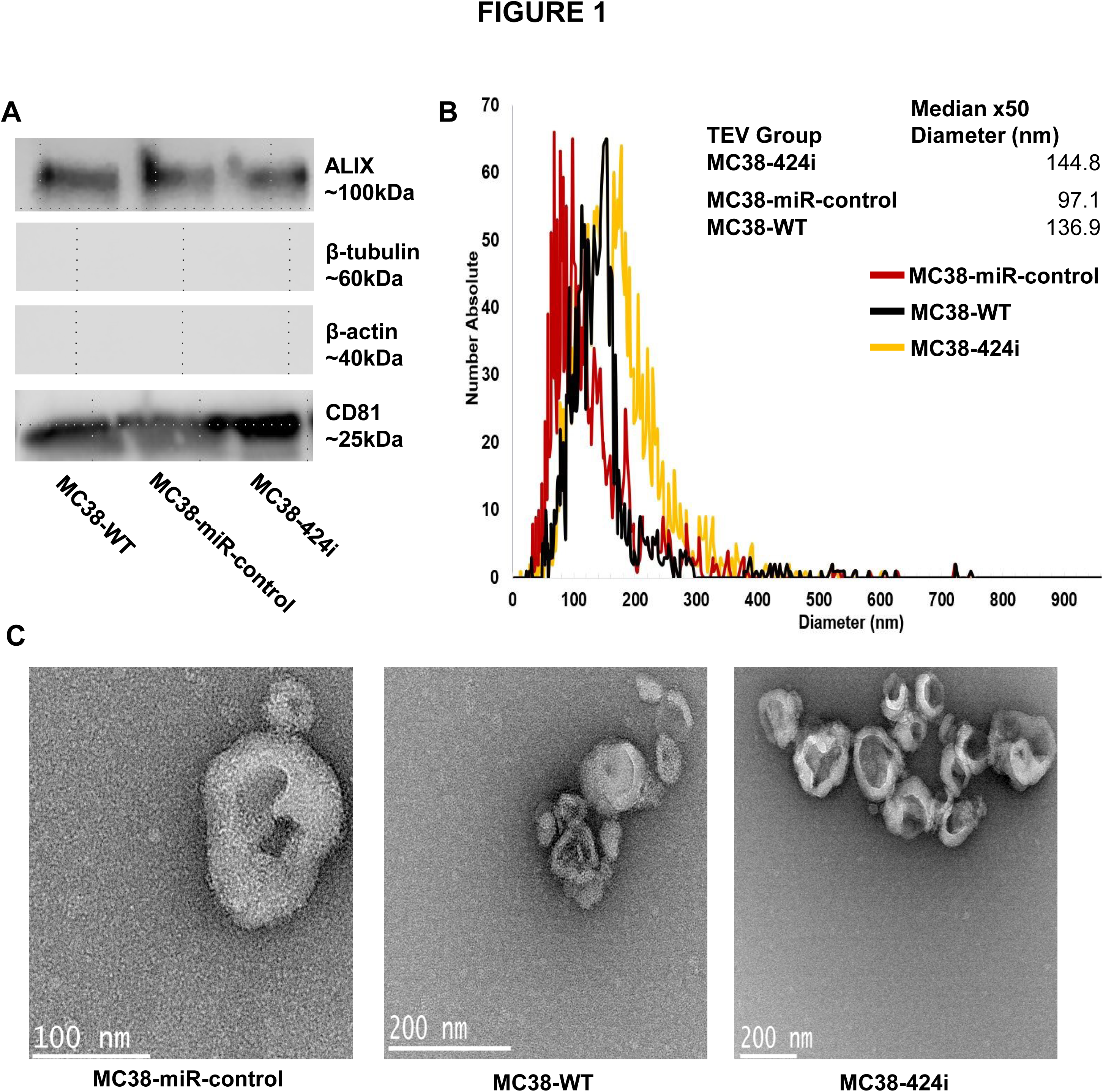
TEV isolation and characterization: **A**) Western Blot data for EV markers ALIX (∼100kDa) and CD81 (∼25kDa) and negative controls β-tubulin (∼60kDa) and β-actin (∼40kDa) for MC38 WT, MC38-miR-control, MC38-424i TEVs. **B)** Nanotracker analysis, size distributions of TEV from MC38-424i (yellow), MC38-miR-control (red), and MC38 WT (black) TEVs. **C)** Transition electron microscopy images of MC38 WT, MC38-miR-control, and MC38-424i TEVs

To test whether allogeneic modified TEVs influenced tumor growth, we prophylactically treated Balb/c mice with 2 doses of 10µg/injection of MC38 WT (n=7), MC38-424i (n=9), MC38miR-control TEVs (n=7), or saline (n=8) **Figure 2A**. We allowed 10 days for an adaptive immune response to develop and challenged mice with CT26 colon cancer cells 2×10^5^ cells/injection and allowed tumors to progress for 21 days. Mice receiving allogeneic MC38-424i TEVs had significantly smaller tumors at the experiment endpoint (71.49 ± 30.32mm^3^) than saline (380.9 ± 95.48mm^3^), allogeneic MC38 WT TEV (284.7 ± 108.1mm^3^) and allogeneic MC38-miR-control TEV (352.3 ± 128.5mm^3^) (p<0.005). Three animals in the MC38-424i TEV group were tumor free following CT26 tumor challenge **Figures 2A and 2C**. Next, we were curious if the observed tumor phenotype was dependent on the presence of CD4 or CD8 T cells. To test, we used 2 *i.p.* injections of CD4 and CD8 (400µg/injection) of depletion antibodies prior to administration of allogeneic TEVs. We confirmed the successful depletion of CD4 and CD8 using flow cytometry comparing depleted mice to naïve spleen and lymph nodes 3 days and 10 days following CD4 and CD8 depletion **Figure 2B**. Following the experimental schema represented in **Figure 2A**, we determined that depletion of CD4 (n=4) and CD8 (n=4) T cells in Balb/c mice was deleterious to the phenotypic effect of the allogeneic modified TEVs on CT26 tumors MC38-424i TEV + anti-CD4 (791.5 ± 85.99mm^3^) MC38-424i TEV + anti-CD8 (619.5 ± 151.5mm^3^) **Figure 2C**. We concluded that MC38-424i TEVs significantly blunted tumor growth compared to MC38 WT and MC38-miR-control TEVs, and saline. Further, the observed phenotype appeared to depend on CD4 and CD8 T cells *in vivo*.

**Figure 2).**
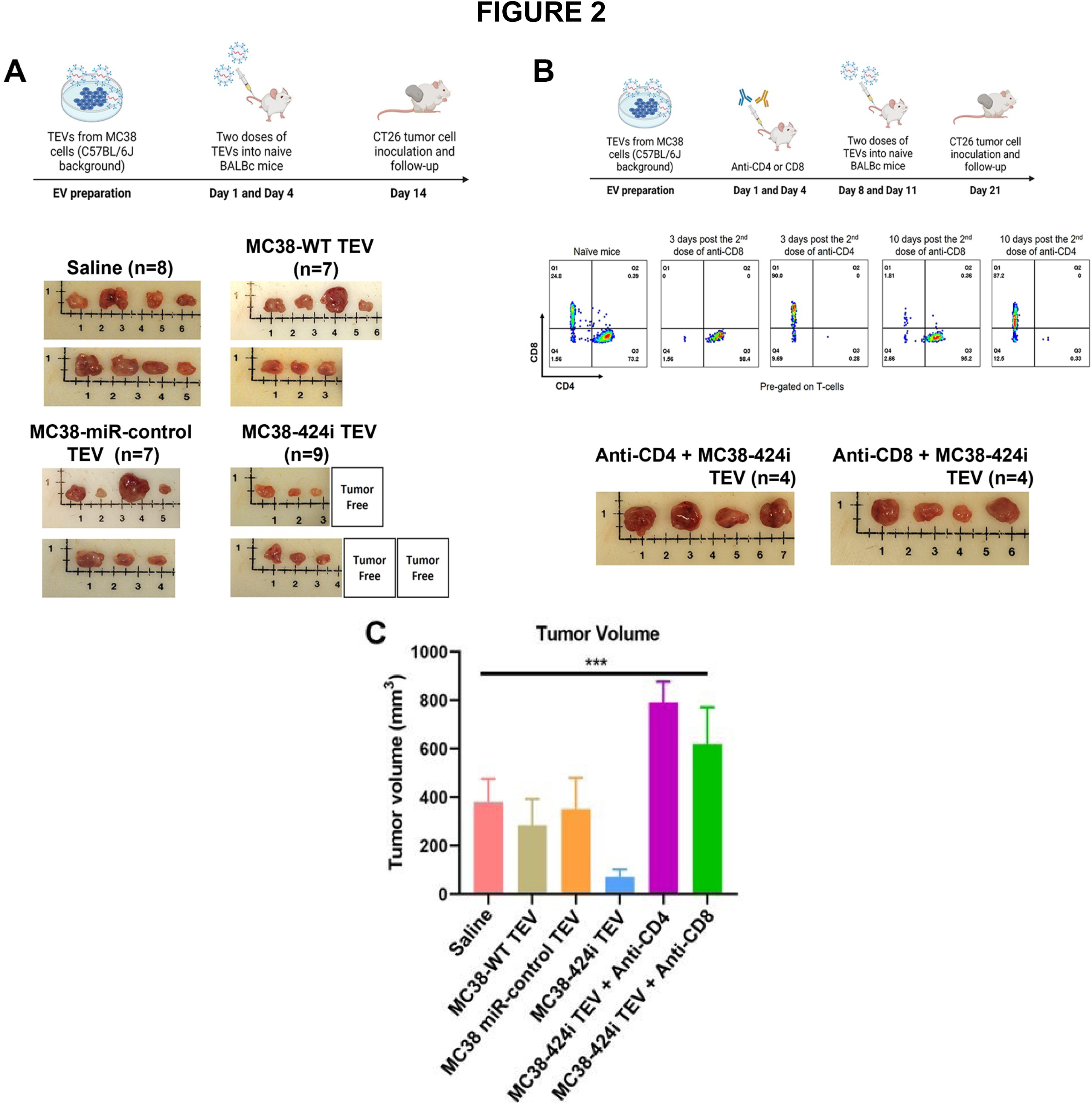
Treatment with allogeneic MC38 TEVs on CT26 tumor slows tumor progression: **A**) Schema of TEV dosing, tumor inoculation, and tumor images within (Saline n=8, MC38 WT TEV n=7, MC38-miR-control TEV n=7, MC38-424i TEV n=9) groups. **B)** α-CD4 and α-CD8 depletion flow cytometry validation and tumor images following TEV inoculation in Balb/c CT26 tumor-bearing mice. **C)** Endpoint tumor volumes between (saline, MC38 WT TEV, MC38 miRcontrol TEV, MC38-424i TEV, MC38-424i TEV + α-CD4, MC38-424i TEV + α-CD8) groups respectively. (*** p<0.005) (Error bars +/-SEM)

### Treatment with Allogeneic Modified TEVs Increases Tumor-infiltrating T cells

After observing a significant effect on tumor growth in the allogeneic MC38-424i TEVs compared to MC38-WT TEVs, MC38-miR-control TEVs, and saline groups, we performed immunofluorescence on tumor tissues to determine if there was a difference in T cell infiltrates in CT26 tumors treated with allogeneic MC38-424i TEVs compared to saline. We observed a significant difference (p<0.05) in CD8^+^ cell count per field (33.11 ± 7.21) and (17.17 ± 3.42) between allogeneic MC38-424i TEVs and saline, respectively **Figure 3A**. Furthermore, given that TEVs can broadly affect cellular responses ^28^, and MC38-424i TEVs are from different genetic background, we investigated the safety profile of MC38-424i TEVs compared to saline. We determined the levels of forty cytokines, such as TNF-alpha, IL-6, IL-2, IL-10, and IFN-gamma, in peripheral blood using a protein array. We didn’t observe positive signals in both groups for some cytokines. Notably, we observed no significant increases in detected cytokines between the allogeneic MC38-424i TEV and saline groups **Figure 3B**. These data suggest that the allogeneic MC38-424i TEVs were associated with tumor T-cell infiltration compared to saline and had a tolerable safety profile.

**Figure 3).**
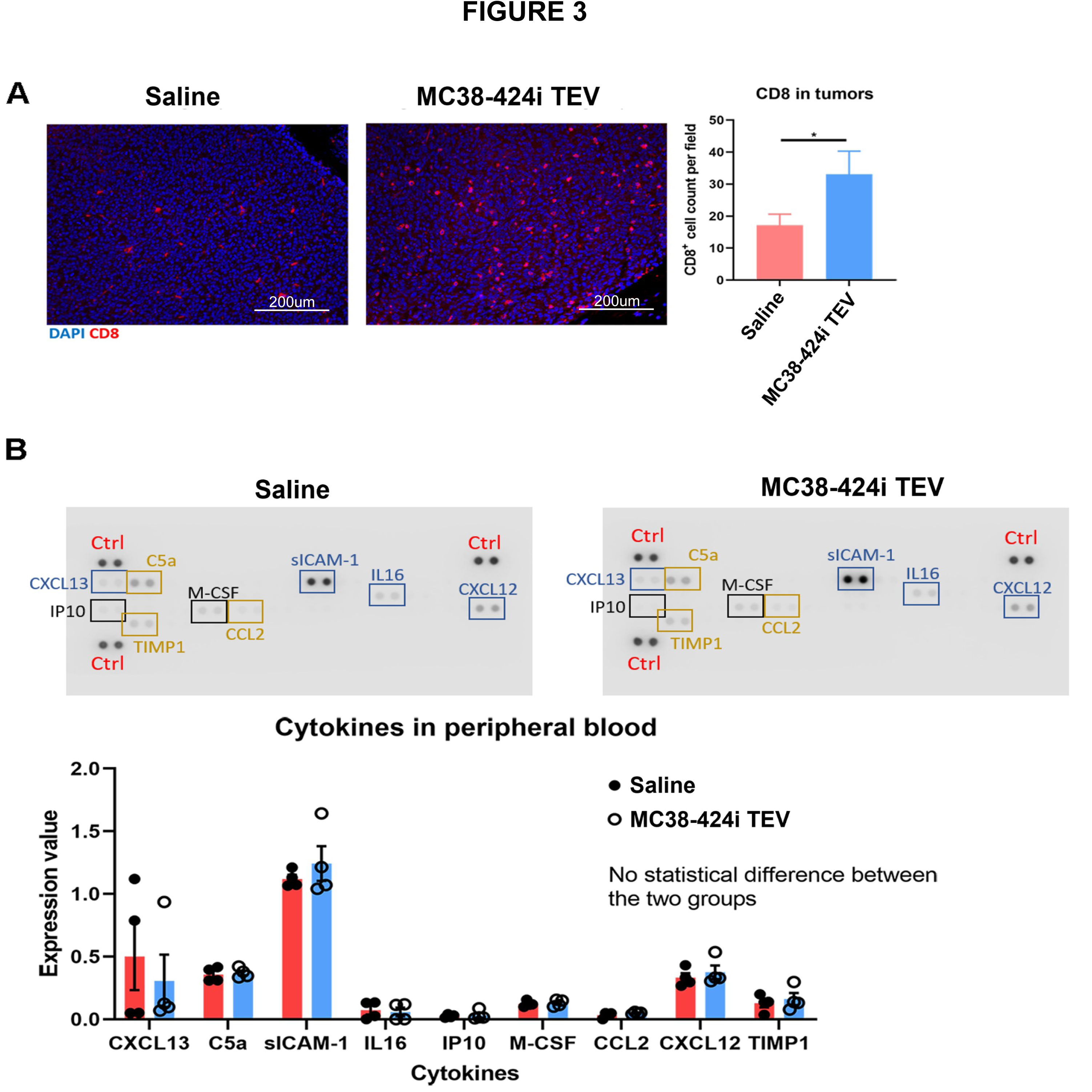
Treatment with allogeneic MC38 TEVs modulates T cell infiltrates: **A**) Immunofluorescence of CD8^+^ T cells (red) DAPI (blue) in CT26 tumors between saline (red) and MC38-424i TEVs (blue). (* p<0.05) (Error bars +/-SEM) **B)** Cytokines in peripheral blood between saline (red) and MC38-424i TEVs (blue) (Error bars +/-SEM).

### Allogeneic Modified TEVs Do Not Significantly Influence B16-F10 Tumor Growth

Having observed a phenotypic effect of MC38 allogeneic MC38-424i TEVs on CT26 colon cancer growth and T cell infiltrate, we were probing if this phenotype was specific to the tissue of the tumor type. To test this, we employed the same prophylactic model using MC38-424i TEVs in C57BL/6 mice that would instead be challenged with B16-F10 melanoma cells **Figure 4A**. Mice received 2 prophylactic doses (10µg/injection) of allogeneic MC38-424i TEVs (n=4) or saline (n=4) on day 1 and day 4. On day 10, C57BL/6 were challenged with subcutaneous injection of 2×10^5^ B16-F10 melanoma cells. We did not observe a significant change in endpoint tumor volumes (1386 ± 536.1mm^3^) and (1657 ± 187.5mm^3^) between allogeneic modified TEV and saline groups **Figure 4B**. Significant tumor volume variability was within the allogeneic MC38-424i TEV group. Furthermore, we did not observe significance in CD8^+^ cell count per field (12.8 ± 3.1) and (15.6 ± 3.3) (p=0.2041) between the allogeneic MC38-424i TEV and saline groups, respectively **Figure 4C**.

**Figure 4).**
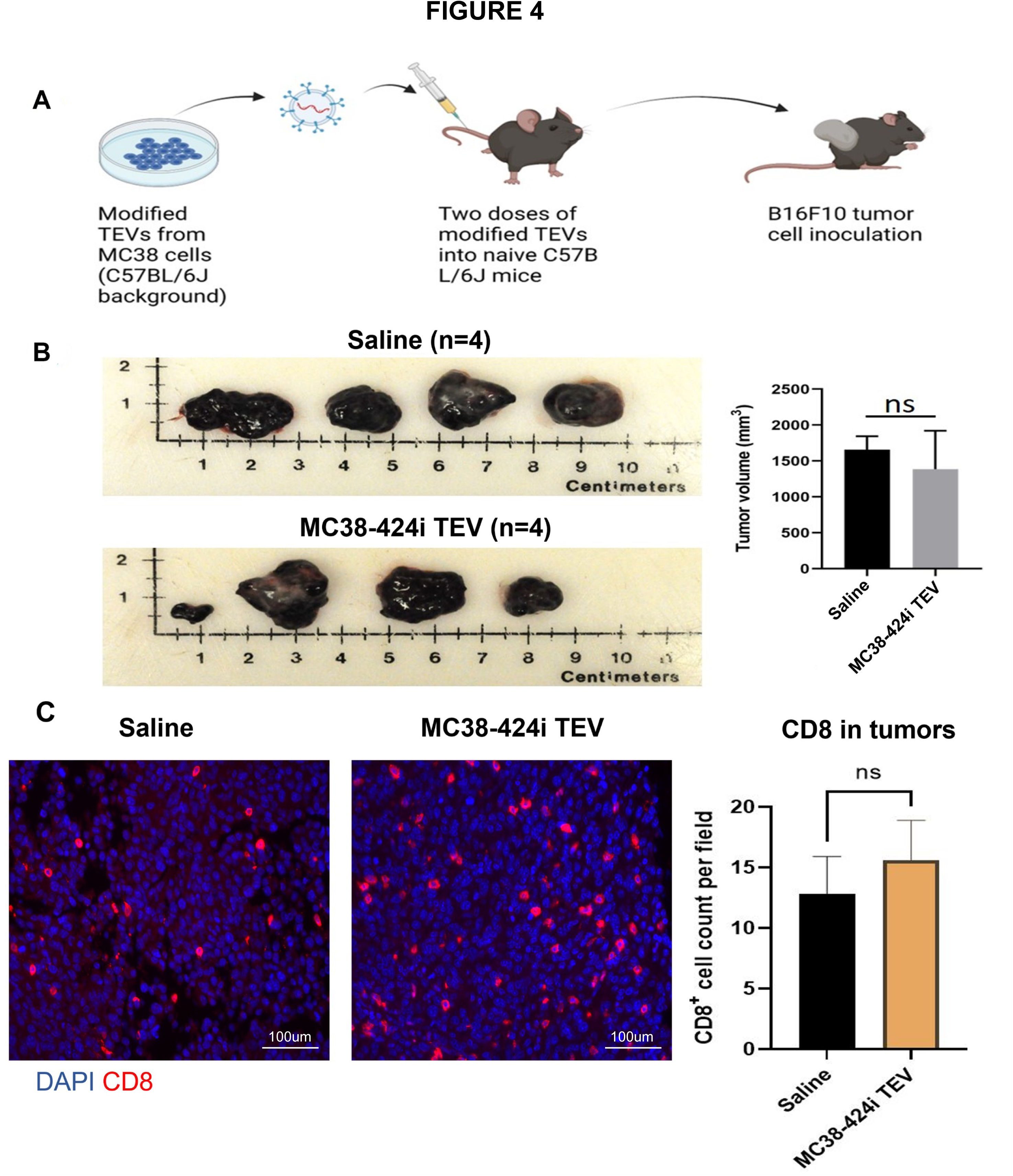
Treatment with MC38 TEVs on B16 melanoma tumors: **A**) MC38 TEV schema in C57BL/6J B16-F10 tumor-bearing animal models. **B)** Tumor volumes between saline (black) and MC38-424i (grey) TEV treatment groups. (ns p>0.05) (Error bars +/-SEM) **C)** Immunofluorescence of CD8^+^ T cells (red) DAPI (blue) in B16-F10 tumors between saline (black) and MC38-424i (orange) TEVs. (ns p>0.05) (Error bars +/-SEM)

### Allogeneic TEVs pulsed dendritic cells are instrumental in stimulating an anti-tumor immune response

Having observed tumor-specific immune activation of allogeneic modified TEVs, we investigate how TEVs are processed *in vitro*. We postulated that TEVs could be captured and presented by dendritic cells (DCs), and tumor antigens within the TEVs can be presented to T cells, stimulating an anti-tumor immune response. This would also explain how the depletion of CD4 and CD8 T cells was indirectly deleterious to the observed phenotype. To observe if TEVs could be captured by DCs, we harvested monocytes and differentiated them into DCs *in vitro* with the addition of IL4, GM-CSF, TNFα, and LPS. We imaged DC populations through Day 6 with or without differentiation factors **Figure 5A**.

**Figure 5).**
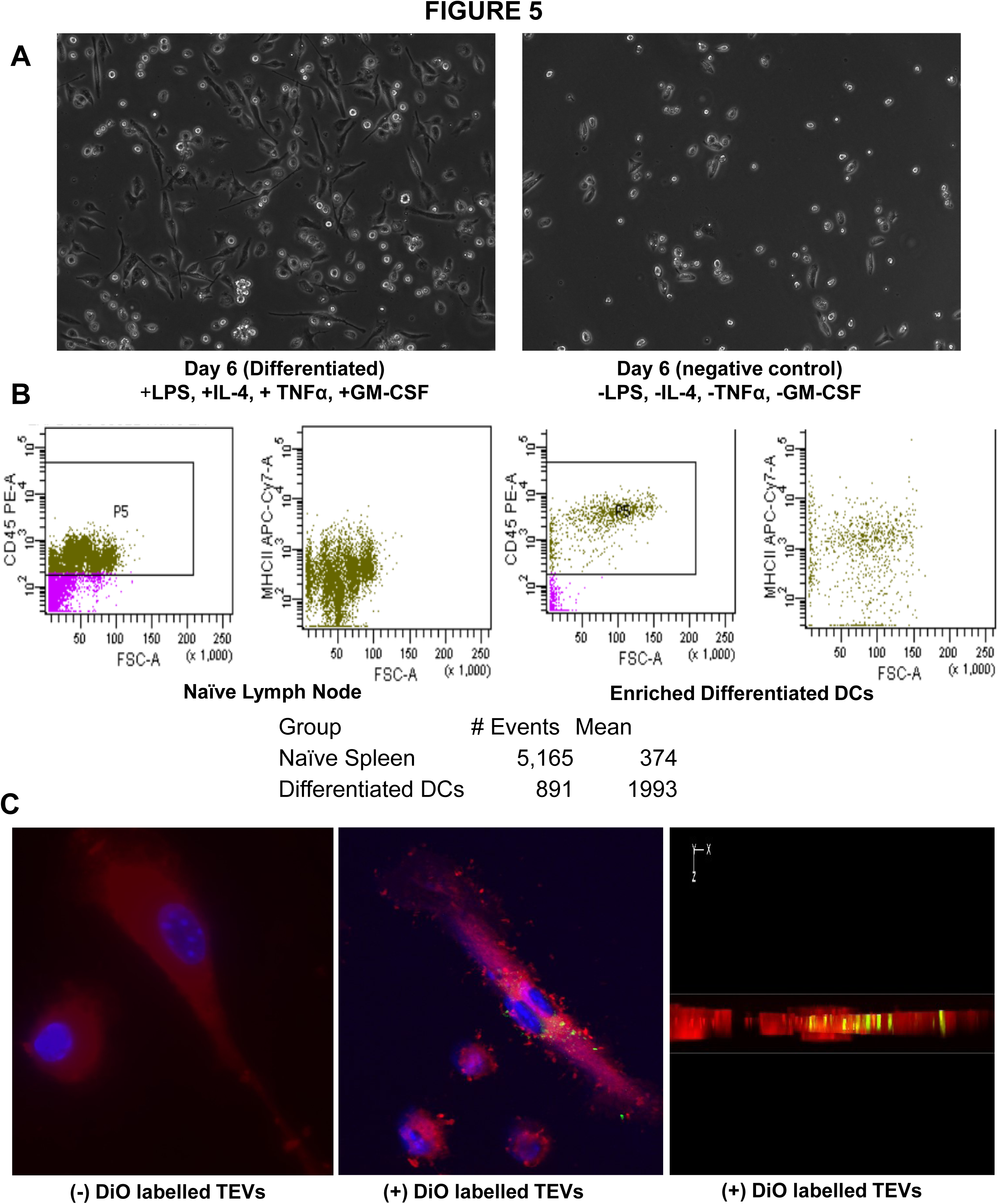
Dendritic cell isolation and TEV capture *in vitro*: **A**) Dendritic cell morphology on day 6 with (left) and without (right) differentiation with IL4, TNFα, GM-CSF, and LPS. **B)** MHCII-APC-Cy7 mean fluorescence intensity of naïve splenocytes and DCs on day 6 *in vitro* differentiation. **C)** Fluorescence microscopy of TEV uptake by DCs (red) with and without DiO-labeled (green) TEVs, DAPI (blue). XZ-plane of Z stack images with DiO TEVs is also presented.

Additionally, we performed flow cytometry and confirmed MHCII expression between enriched DCs (1993 mean signal intensity) and undifferentiated monocytes from naïve spleen and lymph nodes (374 mean signal intensity) **Figure 5B**. We then performed an *in vitro* TEV uptake experiment. We stained the TEVs with the lipophilic dye DiO and exposed the stained TEVs to DCs cultured on glass slides coated with fibronectin (5ug/mL). The stained TEVs were incubated with the DC culture for 24 hours. We observed intracellular localization of the DiO signal (green) within the X-Z plane of the cytopainter (red) stained DCs **Figure 5C**.

Knowing that DCs could capture MC38 TEVs, we wanted to test if allogeneic MC38-424i TEVs pulsed DCs from a Balb/c background in culture could be protective against CT26 tumor challenge. This strategy would allow the delivery of TEVs without the administration of TEVs directly into the bloodstream. To test, we enriched DCs from the spleen of Balb/c mice and exposed them to MC38 WT, Control, and 424i TEVs on day 6 of DC differentiation. The following day we autologously transferred 1×10^6^ DCs exposed to TEVs through *i.v.* tail vein injection. We used 5 groups of Balb/c mice for this study (n=5/group) MC38 WT TEV, MC38-424i TEV, MC38-miR-control-TEV, -No TEV, and saline. We waited 14 days after DC administration before the CT26 tumor challenge and then allowed tumors to progress for 21 days **Figure 6A**. We observed a significant difference in tumor volumes between all TEV groups compared to DCs without TEVs (1105 ± 37.4mm^3^) (n=5) and saline (1186.4 ± 25.2mm^3^) (n=5) groups (p<0.05). However, we did not observe a significant difference in tumor volumes between MC38 WT TEV (834.4 ± 65.4mm^3^) (n=5), MC38-miR-control TEV (910.6 ± 84.4mm^3^) (n=5), and MC38-424i TEV (655.8 ± 355.4mm^3^) (n=5) groups **Figure 6B and 6C**. We did observe a significant difference in tumor volumes when comparing MC38-424i TEVs (655.8 ± 355.4mm^3^) when comparing to No TEVs (1105 ± 37.4mm^3^) and saline groups (1186.4 ± 25.2mm^3^) (p=0.028). Even though we did not observe a significant difference in tumor volumes between TEV groups, we still interrogated the CD8^+^ T cell infiltrates. We observed a significant difference and CD8^+^ (29.4 ± 8.6), (18.0 ± 5.2), and (16.4 ± 4.0) cells per field between the MC38-424i TEV, MC38 WT TEV, and saline groups, respectively (p<0.05) between the MC38-424i TEV, MC38 WT TEV, and saline groups **Figure 6D**.

**Figure 6).**
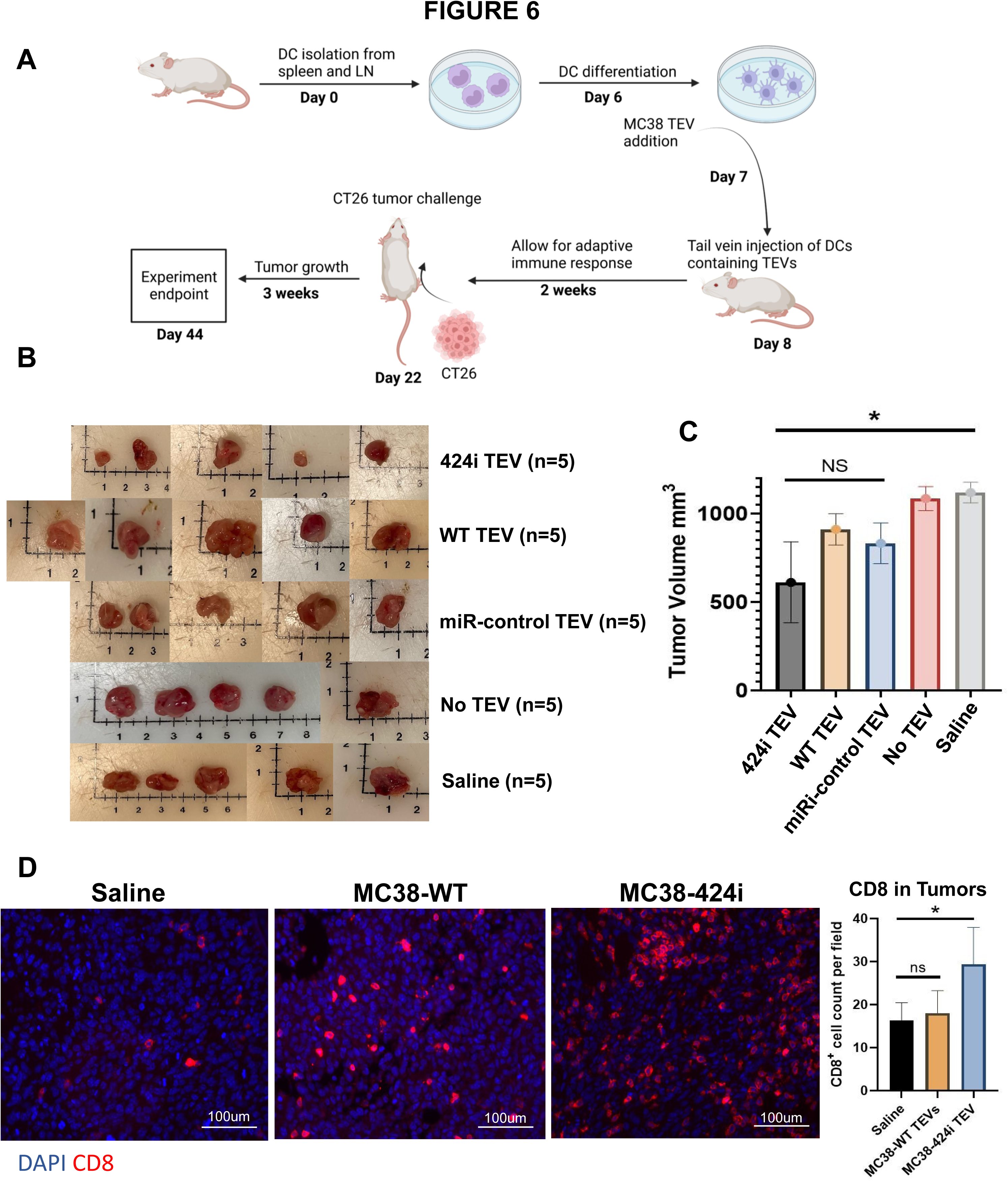
Autologous transfer of DCs exposed to TEVs slows tumor growth: **A**) Schema of DC isolation, TEV pulsing, and autologous transfer of DCs to Balb/c animals prior to tumor challenge with CT26 colon cancer cells. **B)** Endpoint tumor images of MC38-424i, MC38 WT, MC38-miR-control, No TEV, and saline groups (n=5/group). **C)** Endpoint tumor volumes of MC38-424i, MC38 WT, MC38-miR-control, No TEV, and saline groups (n=5/group). (* p<0.05) (ns p>0.05) (Error bars +/-SEM). **D)** Immunofluorescence and counts per field of CD8^+^ T cells (red) DAPI (blue) in CT26 tumors between saline, MC38WT, and MC38-424i TEVs. (* p<0.05) (ns p>0.05) (Error bars +/-SEM).

## DISCUSSION

Here we sought to investigate the consequences of the administration of MC38 allogeneic TEVs lacking functional miR-424 on tumor growth and immune infiltrates of mice bearing CT26 and B16-F10 tumors. From our prior investigations, we hypothesized that MC38-424i TEVs would be capable of suppressing tumor growth of CT26 and B16-F10 tumors.^19^ We observed a significant reduction in tumor volumes when MC38-424i TEVs were prophylactically administered to mice challenged with CT26 tumors **Figure 2A and 2C**. However, we did not observe any significant effects on endpoint tumor volumes when dosing C57BL/6 animals with MC38-424i TEVs when challenged with B16-F10 tumors **Figure 4B**. Yet, it is important to note that two animals did observe smaller tumors at the endpoint, but within-group variability did not yield statistical significance. It has been reported that there are conserved tumor exon junctions between MC38 colon cancer cells B16-F10 melanoma cells that can be presented on MHCI.^29^ Additionally, mice exposed prophylactically to conserved tumor exon junctions showed protective effects against B16-F10 growth through induction of anti-tumor immune responses.^29^ We speculate that conserved tumor neoantigens between MC38 TEVs and B16-F10 melanoma may undergo immunoediting, thus allowing for immune escape.^30, 31^

Conversely, the tumor neoantigens conserved between MC38 TEVs and B16-F10 may only stimulate a weak immune response.^32^ At the same time, it has been reported that MC38 cells and CT26 cell lines have similar somatic mutations per megabase resulting in a similar tumor mutational burden.^33^ This could explain why a more robust immune response **(Figure 3A)** and protective effects **(Figure 2C)** on tumor growth were observed in our study with MC38-424i TEVs in CT26 tumors but not as robust in B16-F10 tumors. Similarly, we observed a significant increase in CD8^+^T cells in CT26 tumors treated with miR-424i TEVs **(Figure 3A),** but did not observe any significant increase in CD8^+^T cells in B16-F10 tumors treated with miR-424i TEVs **Figure 4C**. These data suggest that tissue of origin or similarities in tumor mutational burden can be determinant of the efficacy of allogeneic TEVs, as the B16-F10 immune infiltrates and tumor growth were not as heavily influenced by the MC38-424i TEVs. Furthermore, we did not notice any significant increases in peripheral blood cytokines when comparing allogeneic MC38-424i TEVs to saline, suggesting that TEVs were relatively well tolerated in an allogeneic context **Figure 3B**. This was critical as any induced cytokine release syndrome profile would render allogeneic TEVs clinically untranslatable.^34, 35^

We also sought to show that TEVs are captured by dendritic cells (DCs) and if DCs that were differentiated *in vitro* could be pulsed with allogeneic MC38 TEVs and could be autologously transferred back into Balb/c mice and initiate an anti-tumor immune response against CT26 tumors. This approach would allow autologous delivery of tumor antigens without directly administering TEVs into the bloodstream. It has been reported that antigen presenting cells (APCs) such as DCs are mediators of effective anti-tumor through presentation of antigens to T cells.^36^ Previous studies have attempted to expose tumor cells and tumor antigens to DCs (Sipluecel-T) and shown various efficacy levels in prostate cancer.^26, 37, 38^ We postulated that the abundance of endogenous TEVs containing miR-424, which we showed in our prior investigation^19^, could still be limiting the effective translational of DC vaccine approaches into clinical implementation. Our data suggest allogeneic TEVs can be pulsed to DCs *in vitro* **Figure 5C**. These DCs can then be autologously transferred to Balb/c mice and can be protective compared to No TEVs and saline **Figures 6A, 6B, and 6C**. Even though we did not observe significant differences between tumor volumes within the allogeneic MC38 TEV groups, we did observe a significant difference in CD8^+^ T cell infiltration in DCs that were pulsed with allogeneic MC38-424i TEVs compared to MC38 WT. Oftentimes, advanced tumors are intrinsically immunosuppressive and unresponsive to ICIs due to a lack of functional CD8^+^ T cell infiltration by various mechanisms.^39, 40^ Our data suggests that DCs pulsed with allogeneic MC38-424i TEVs could promote tumor CD8^+^ T cell infiltration **Figure 6D**. Additionally, MC38-424i TEVs pulsed to DCs can promote anti-tumor immune responses without directly administering TEVs into the blood of Balb/c mice **Figure 5C**. Future investigations should combine autologous transfer of DCs pulsed with MC38-424i TEVs to determine if ICI efficacy is influenced in CRC preclinical models before clinical translation.

## CONCLUSION

In this study, we observed that allogeneic MC38 TEVs lacking functional miR-424, when treated prophylactically, can be protective against CT26 tumor growth through the induction of anti-tumor immune responses. We also showed that allogeneic MC38-424i TEVs were well tolerated and did not significantly alter the expression of cytokines in peripheral blood. Furthermore, we showed that allogeneic MC38 TEVs could be pulsed to DCs. When autologously transferred to mice was protective against CT26 tumor challenge and resulted in increased CD8^+^T cell infiltrates compared to MC38 WT TEVs and saline. These data together suggest that further investigation is warranted as to how modified TEVs pulsed to DCs can be combined with ICIs in additional preclinical models of CRC to see how ICI efficacy is altered in the presence of DCs pulsed with allogeneic TEVs. Experiments coupling combinations of immunotherapies designed to stimulate DCs and ICIs will be essential to conduct prior to clinical translation and implementation.

## ACKNOWLEDGMENTS

We thank the Masonic Cancer Center’s core facilities and the Clinical and translational sciences institute for supporting histological services. Biorender.com was used to generate experimental schematics. This study is supported by research grants from the Minnesota Colorectal Cancer Funds, Mezin Koat colorectal cancer research fund, and research funds from the Department of Surgery and CTSI, University of Minnesota. The Minnesota Colorectal Cancer Research Foundation supports TJG and XZ graduate fellowships.

## ETHICS STATEMENT

This study followed the University of Minnesota Institutional Animal Care and Use Committee protocol code 2107-39273A 11-02-21.

## AUTHOR CONTRIBUTIONS

All authors provided meaningful intellectual contributions to this work and were involved in its preparation and editing. TJG, XZ, and SS conceived the study. TJG, XZ, DW, and SS generated and analyzed the data. TJG, XZ, and SS wrote the manuscript, and all the authors edited the manuscript.

## DATA AVAILABILITY

No large datasets was generated or analyzed in this study.

